# Factors influencing leaf- and root-associated communities of bacteria and fungi across 33 plant orders in a grassland

**DOI:** 10.1101/439646

**Authors:** Hirokazu Toju, Hiroko Kurokawa, Tanaka Kenta

## Abstract

In terrestrial ecosystems, plants interact with diverse taxonomic groups of bacteria and fungi in the phyllosphere and rhizosphere. Although recent studies based on high-throughput DNA sequencing have drastically increased our understanding of plant-associated microbiomes, we still have limited knowledge of how plant species in a species-rich community differ in their leaf and root microbiome compositions. In a cool-temperate semi-natural grassland in Japan, we compared leaf- and root-associated microbiomes across 138 plant species belonging to 33 plant orders. Based on the whole-microbiome inventory data, we analyzed how sampling season as well as the taxonomy, nativeness (native or alien), lifeform (herbaceous or woody), and mycorrhizal type of host plants could contribute to variation in microbiome compositions among co-occurring plant species. The data also allowed us to explore prokaryote and fungal lineages showing preferences for specific host characteristics. The list of microbial taxa showing significant host preferences involved those potentially having some impacts on survival, growth, or environmental resistance of host plants. Overall, this study provides a platform for understanding how plant and microbial communities are linked with each other at the ecosystem level.

## INTRODUCTION

Plants interact with various taxonomic groups of microbes both in the phyllosphere and rhizosphere (van der Heijden et al., 1998;Berendsen et al., 2012;Bai et al., 2015;Peay et al., 2016). Diverse bacteria and yeasts, for example, are present on leaf surfaces, involved in underappreciated metabolic pathways (Mercier and Lindow, 2000;Delmotte et al., 2009;Hacquard et al., 2015). In addition to those epiphytes, a number of bacteria and filamentous fungi are known to inhabit leaf tissue, potentially playing pivotal roles in resistance of host plants against biotic and abiotic environmental stresses (Schardl and Phillips, 1997;Arnold et al., 2003). In root systems, mycorrhizal fungi provide plants with soil phosphorus and/or nitrogen, fueling hosts’ growth (Parniske, 2008;Smith and Read, 2008;Tedersoo et al., 2010). Likewise, some endophytic fungal taxa have known to convert organic forms of nitrogen into inorganic forms, contributing to physiological conditions of host plants (Newsham, 2011). Moreover, endophytic bacteria and fungi associated with roots can increase disease resistance of host plants, possibly by stimulating host immune systems (Ramamoorthy et al., 2001;Pieterse et al., 2014;Hacquard et al., 2017) or by suppressing soil pathogens with antibiotic chemicals (Compant et al., 2005;Gao et al., 2010). Thus, understanding of the compositions of plant microbiomes is a prerequisite for understanding the physiology and ecology of plants in terrestrial ecosystems (van der Heijden et al., 2008;Schlaeppi and Bulgarelli, 2015;Toju et al., 2018a).

While exploration of plant microbiomes has been accelerated since the emergence of high-throughput DNA sequencing (Öpik et al., 2009;Lundberg et al., 2012;Bai et al., 2015), we still have limited knowledge of how diverse plant species co-occurring in a grassland or forest ecosystem can differ in their microbiome compositions (Toju et al., 2016a). Moreover, most plant microbiome studies target only bacteria or fungi [but see (Agler et al., 2016)) in either above-or below-ground systems [but see (Bai et al., 2015;Wagner et al., 2016)], precluding comprehensive understanding of microbiome compositions. Given that bacteria and fungi can interact with each other within hosts (Frey-Klett et al., 2007;Hoffman and Arnold, 2010) and that above- and below-ground ecological processes can be interlinked (Bever et al., 2010;Mangan et al., 2010;Van der Putten et al., 2013), the targets of plant microbiome studies need to be expanded towards a better understanding of the processes that organize plant and microbial communities in the wild. Studies comparing microbiome compositions across tens (or more) of plant species co-occurring in natural ecosystems (Toju et al., 2014;Toju et al., 2018b), in particular, will allow us to examine what kinds of host properties can contribute to the organization of leaf- and root-associated microbial communities.

In this study, we sampled leaves and roots of 138 plant species representing 112 genera, 55 families, and 33 orders in a cool-temperate grassland in Japan, thereby performing a high-throughput sequencing analysis of both prokaryote and fungal communities associated with plants. The sample set of diverse plant species allowed us to examine what host properties can contribute to variation in leaf and root microbiome compositions in an ecosystem. Furthermore, we statistically tested how each prokaryote or fungal genus showed preferences for seasons as well as preference for nativeness (native or alien), lifeform (herbaceous or woody), and mycorrhizal type (ectomycorrhizal, arbuscular mycorrhizal, non-mycorrhizal, or variable mycorrhizal) of host plants. Overall, this study, for the first time, shows how more than 100 plant species in a single ecosystem can differ in their leaf and root microbiome compositions depending on their characteristics. The statistical results on plant–microbe associations shed light on underappreciated diversity of host–symbiont associations in grasslands, providing fundamental information for conserving and restoring terrestrial ecosystems.

## MATERIALS AND METHODS

### Sampling

Fieldwork was conducted in Sugadaira Research Station, Mountain Science Center, University of Tsukuba, Sugadaira, Ueda, Nagano Prefecture, Japan (36.524 °N; 138.349 °E; 1340 m asl). In Sugadaira Research Station, 6 ha of a semi-natural grassland has been maintained by mowing plants in autumn and thereby preventing the community succession to a forest. Thus, woody plant species that occurred in the grassland are shrubs or saplings of tall trees colonized from surrounding forests. In total, 200 plant species have been reported from the grassland, including some alien species.

In the grassland, both native and alien plant species were sampled to reveal the compositions of prokaryote and fungal communities associated with leaves and roots through summer and autumn (July 19-20, August 16-18, and September 7-8) in 2017. We targeted only non-reproductive plant individuals that had neither flower buds, flowers, nor fruits so that plant physiology and chemistry would not be affected by reproduction. We tried to sample as many plant species as possible within the sampling days in each month. Note that root systems of multiple plant species were tangled with each other at the study site due to the dominance of perennial plants. Therefore, we sampled 1–8 liters of soil including root systems for each target plant individuals and quickly washed the root system in a nearby laboratory to carefully trace root tips directly connected to above-ground tissue of the target plant. A 1-cm^2^ disc of a mature leaf and a 2-cm segment of a terminal root were collected from each plant sample and preserved at −20 °C until DNA extraction. After the sampling, remaining plant organs of rare plant species were replaced at the original sampling positions. In total, 289 plant individuals representing 138 plant species (112 genera, 55 families, 33 orders) were collected (Supplementary Data 1). The permission of sampling was issued by Sugadaira Research Station, Mountain Science Center, University of Tsukuba.

### DNA Extraction, PCR, and Sequencing

Each leaf or root sample was surface-sterilized by immersing it in ×1/100 NaClO (Nacalai Tesque) for 1 min and it was subsequently washed in ethanol twice. DNA extraction was extracted with a cetyltrimethylammonium bromide (CTAB) method after pulverizing the roots with 4 mm zirconium balls at 25 Hz for 3 min using a TissueLyser II (Qiagen) (Toju et al., 2013).

For each of the leaf and root samples, the 16S rRNA V4 region of the prokaryotes and the internal transcribed spacer 1 (ITS1) region of fungi were amplified. The PCR of the 16S rRNA region was performed with the forward primer 515f (Caporaso et al., 2011) fused with 3–6-mer Ns for improved Illumina sequencing quality (Lundberg et al., 2013) and the forward Illumina sequencing primer (5’-TCG TCG GCA GCG TCA GAT GTG TAT AAG AGA CAG-[3-6-mer Ns] - [515f]-3’) and the reverse primer 806rB (Apprill et al., 2015) fused with 3–6-mer Ns and the reverse sequencing primer (5’-GTC TCG TGG GCT CGG AGA TGT GTA TAA GAG ACA G [3–6-mer Ns] - [806rB]-3’) (0.2 μM each). To prevent the amplification of mitochondrial and chloroplast 16S rRNA sequences of host plants, specific peptide nucleic acids [mPNA and pPNA; Lundberg et al. (2013)] (0.7 μM each) were added to the reaction mix of KOD FX Neo (Toyobo). The temperature profile of the PCR was 94 °C for 2 min, followed by 35 cycles at 98 °C (denaturation) for 10 s, 78 °C (annealing of PNA) for 10 s, 60 °C (annealing of primers) for 30 s, and 68 °C (extension) for 50 s, and a final extension at 68 °C for 5 min. To prevent generation of chimeric sequences, the ramp rate through the thermal cycles was set to 1 °C/sec (Stevens et al., 2013). Illumina sequencing adaptors were then added to respective samples in the supplemental PCR using the forward fusion primers consisting of the P5 Illumina adaptor, 8-mer indexes for sample identification (Hamady et al., 2008) and a partial sequence of the sequencing primer (5’-AAT GAT ACG GCG ACC ACC GAG ATC TAC AC - [8-mer index] - TCG TCG GCA GCG TC-3’) and the reverse fusion primers consisting of the P7 adaptor, 8-mer indexes, and a partial sequence of the sequencing primer (5’-CAA GCA GAA GAC GGC ATA CGA GAT - [8-mer index] - GTC TCG TGG GCT CGG-3’). KOD FX Neo was used with a temperature profile of 94 °C for 2 min, followed by 8 cycles at 98 °C for 10 s, 55 °C for 30 s, and 68 °C for 50 s (ramp rate = 1 °C/s), and a final extension at 68 °C for 5 min. The PCR amplicons of the samples were then pooled after a purification/equalization process with the AMPureXP Kit (Beckman Coulter). Primer dimers, which were shorter than 200 bp, were removed from the pooled library by supplemental purification with AMpureXP: the ratio of AMPureXP reagent to the pooled library was set to 0.6 (v/v) in this process.

The PCR of fungal ITS1 region was performed with the forward primer ITS1F_KYO1 (Toju et al., 2012) fused with 3–6-mer Ns for improved Illumina sequencing quality (Lundberg et al., 2013) and the forward Illumina sequencing primer (5’-TCG TCG GCA GCG TCA GAT GTG TAT AAG AGA CAG-[3–6-mer Ns] – [ITS1F_KYO1]-3’) and the reverse primer ITS2_KYO2 (Toju et al., 2012) fused with 3–6-mer Ns and the reverse sequencing primer (5’-GTC TCG TGG GCT CGG AGA TGT GTA TAA GAG ACA G [3–6-mer Ns] - [ITS2_KYO2]-3’). The buffer and polymerase system of KOD FX Neo was used with a temperature profile of 94 °C for 2 min, followed by 35 cycles at 98 °C for 10 s, 58 °C for 30 s, and 68 °C for 50 s, and a final extension at 68 °C for 5 min. Illumina sequencing adaptors and 8-mer index sequences were then added in the second PCR as described above. The amplicons were purified and pooled as described above.

The sequencing libraries of the prokaryote 16S and fungal ITS regions were processed in an Illumina MiSeq sequencer (run center: KYOTO-HE; 15% PhiX spike-in). Because the quality of forward sequences is generally higher than that of reverse sequences in Illumina sequencing, we optimized the MiSeq run setting in order to use only forward sequences. Specifically, the run length was set 271 forward (R1) and 31 reverse (R4) cycles to enhance forward sequencing data: the reverse sequences were used only for discriminating between 16S and ITS1 sequences based on the sequences of primer positions.

### Bioinformatics

The raw sequencing data were converted into FASTQ files using the program bcl2fastq 1.8.4 distributed by Illumina. The output FASTQ files were demultiplexed with the program Claident v0.2. 2018.05.29 (Tanabe and Toju, 2013;Tanabe, 2018), by which sequencing reads whose 8-mer index positions included nucleotides with low (< 30) quality scores were removed. Only forward sequences were used in the following analyses after removing low-quality 3’-ends using Claident. Noisy reads (Tanabe, 2018) were subsequently discarded and then denoised dataset consisting of 2,973,811 16S and 2,774,197 ITS1 reads were obtained. The sequencing data were deposited to DNA Data Bank of Japan (DDBJ) (DRA007062).

For each dataset of 16S and ITS1 regions, filtered reads were clustered with a cut-off sequencing similarity of 97% using the program VSEARCH (Rognes et al., 2014) as implemented in Claident. The operational taxonomic units (OTUs) representing less than 10 sequencing reads were subsequently discarded (Supplementary Data 2). The molecular identification of the remaining OTUs was performed based on the combination of the query-centric auto-*k*-nearest neighbor (QCauto) method (Tanabe and Toju, 2013) and the lowest common ancestor (LCA) algorithm (Huson et al., 2007) as implemented in Claident (Supplementary Data 2). Note that taxonomic identification results based on the combination of the QCauto search and the LCA taxonomic assignment are comparable to, or sometimes more accurate than, those with alternative approaches (Tanabe and Toju, 2013;Toju et al., 2016a;Toju et al., 2016b).

For each combination of target region (16S or ITS1) and sample type (root or soil), we obtained a sample × OTU matrix, in which a cell entry depicted the number of sequencing reads of an OTU in a sample (Supplementary Data 3). The cell entries whose read counts represented less than 0.1% of the total read count of each sample were removed to minimize effects of PCR/sequencing errors (Peay et al., 2015). The filtered matrix was then rarefied to 500 reads per sample using the “rrarefy” function of the vegan 2.5-2 package (Oksanen et al., 2012) of R 3.5.1 (R-Core-Team, 2018). Samples with less than 500 reads were discarded in this process: the numbers of OTUs in the rarefied sample × OTU matrices were 1,470, 5,638, 1,537, and 3,367 for leaf prokaryote, root prokaryote, leaf fungal, and root fungal datasets, respectively (Supplementary Data 4). For each dataset, we also obtained order- and genus-level matrices, which represented order- and genus-level taxonomic compositions of microbes (prokaryotes or fungi), respectively (Supplementary Data 5).

### Prokaryote and Fungal Diversity

Relationships between the number of sequencing reads and that of detected OTUs were examined for respective data matrices (leaf prokaryote, root prokaryote, leaf fungal, and root fungal datasets) with the “rarecurve” function of the R vegan package. Likewise, relationships between the number of samples and that of prokaryote/fungal orders or genera were examined with the vegan “specaccum” function. The order-level taxonomic compositions of leaf prokaryotes, root prokaryotes, leaf fungi, and root fungi were visualized in bar graphs for respective plant orders.

### Factors Contributing to Microbiome Compositions

For each dataset (leaf prokaryote, root prokaryote, leaf fungal, or root fungal dataset), factors contributing to microbial community compositions were examined with the permutational analysis of variance [PERMANOVA; Anderson (2001)] using the vegan “adonis” function (10,000 permutations). Sampling month (July, August, or September) and four variables representing host plant properties were included as explanatory variables. Specifically, order-level plant taxonomy, plant nativeness (native or alien), plant lifeform (herbaceous or woody), and plant mycorrhizal type [ectomycorrhizal (EcM), arbuscular mycorrhizal (AM), non-mycorrhizal (NM), or variable mycorrhizal (NM-AM)] (Brundrett, 2009) were included as variables representing host properties. In each model, a matrix representing order-or genus-level taxonomic compositions of prokaryotes/fungi was used as the input response matrix. The “Bray-Curtis” metric of *β*-diversity was used in the PERMANOVA analyses.

### Randomization Analyses of Preferences

To explore prokaryote/fungal genera that preferentially occurred on plant samples with specific properties, a series of randomization tests were performed. In each genus-level matrix (leaf prokaryote, root prokaryote, leaf fungal, or root fungal genus-level matrix), sample information was shuffled among plant samples (100,000 permutations) and then preference of a prokaryote/fungal genus (*i*) for a sample property (*j*) was evaluated as follows:

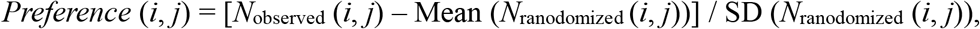

where *N*_observed_ (*i*, *j*) denoted the mean number of the sequencing reads of genus *i* among property *j* samples in the original data, and the Mean (*N*_ranodomized_ (*i*, *j*)) and SD (*N*_ranodomized_ (*i*, *j*)) were the mean and standard deviation of the number of sequencing reads for the focal genus–sample property combination across randomized matrices. Based on the index, preferences for sampling month (July, August, or September), plant nativeness (native or alien), plant lifeform (herbaceous or woody), and plant mycorrhizal type were examined, respectively. Because most plant orders included a few plant species in our datasets, the preference analysis was not applied to plant taxonomy. Regarding this standardized preference index, values larger than three generally represent strong preferences [false discovery rate (FDR) < 0.05; Toju et al. (2016a)]: hence, we listed genera whose preference values exceeded three.

## RESULTS

### Prokaryote and Fungal Diversity

After a series of quality filtering and rarefaction procedures, 41.1 (SD = 22.1), 143.4 (SD = 37.9), 54.5 (SD = 18.8), and 46.0 (SD = 22.5) OTUs per sample, on average, were detected, from the leaf prokaryote, root prokaryote, leaf fungal, and root fungal datasets, respectively (Supplementary Fig. 1). The numbers of prokaryote orders and genera were higher in root samples than in leaf samples, while those of fungal orders and genera showed opposite patterns (Fig. 2).

**FIGURE 1.**
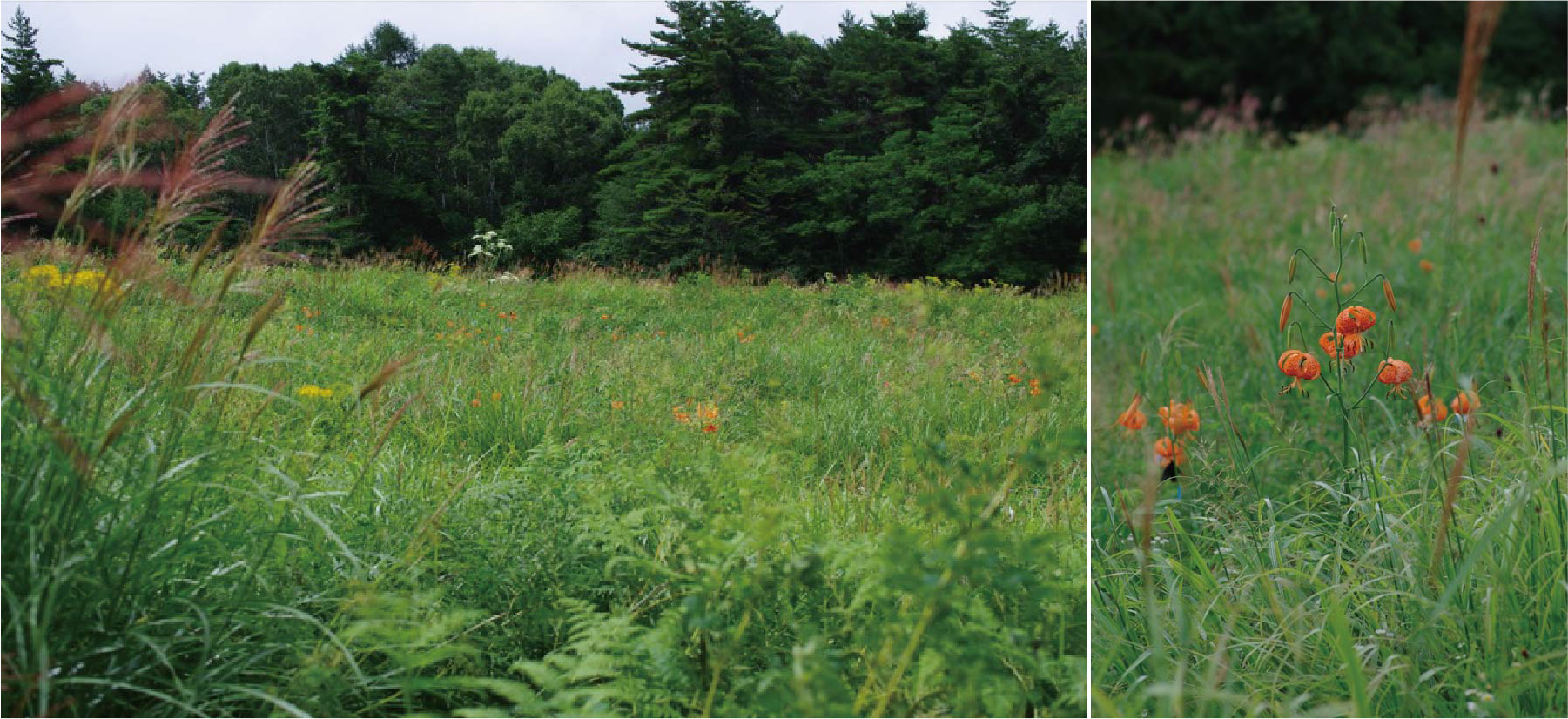
Grassland of Sugadaira Research Station.

**FIGURE 2.**
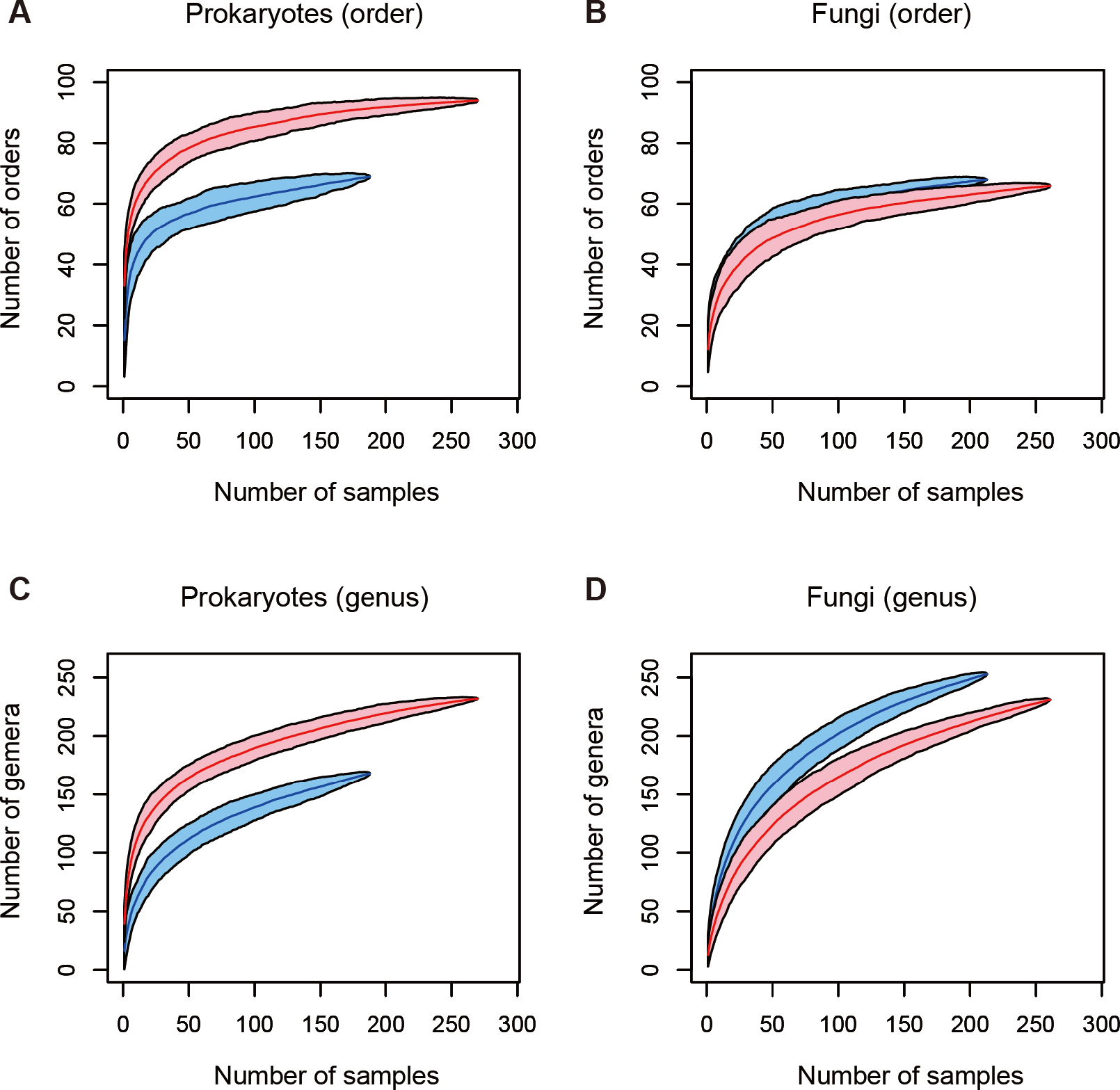
Relationship between the number of leaf/root samples and that of prokaryote/fungal taxa observed. **(A)** Number of prokaryote orders. In total, sequencing data were successfully obtained from 188 leaf and 270 root samples. Blue and red curves represent leaf and root samples, respectively. **(B)** Number of fungal orders. In total, sequencing data were successfully obtained from 213 leaf and 261 root samples. **(C)** Number of prokaryote genera (188 leaf and 270 root samples). **(D)** Number of fungal genera (213 leaf and 261 root samples).

The leaf prokaryote communities of the examined plants were dominated by the order Rhizobiales, while diverse bacterial taxa constituted the root prokaryote communities (Fig. 3A-B). In the leaf fungal communities, the order Capnodiales were the most abundant, while root fungal community compositions varied considerably among host plant orders (Fig. 3C-D).

**FIGURE 3.**
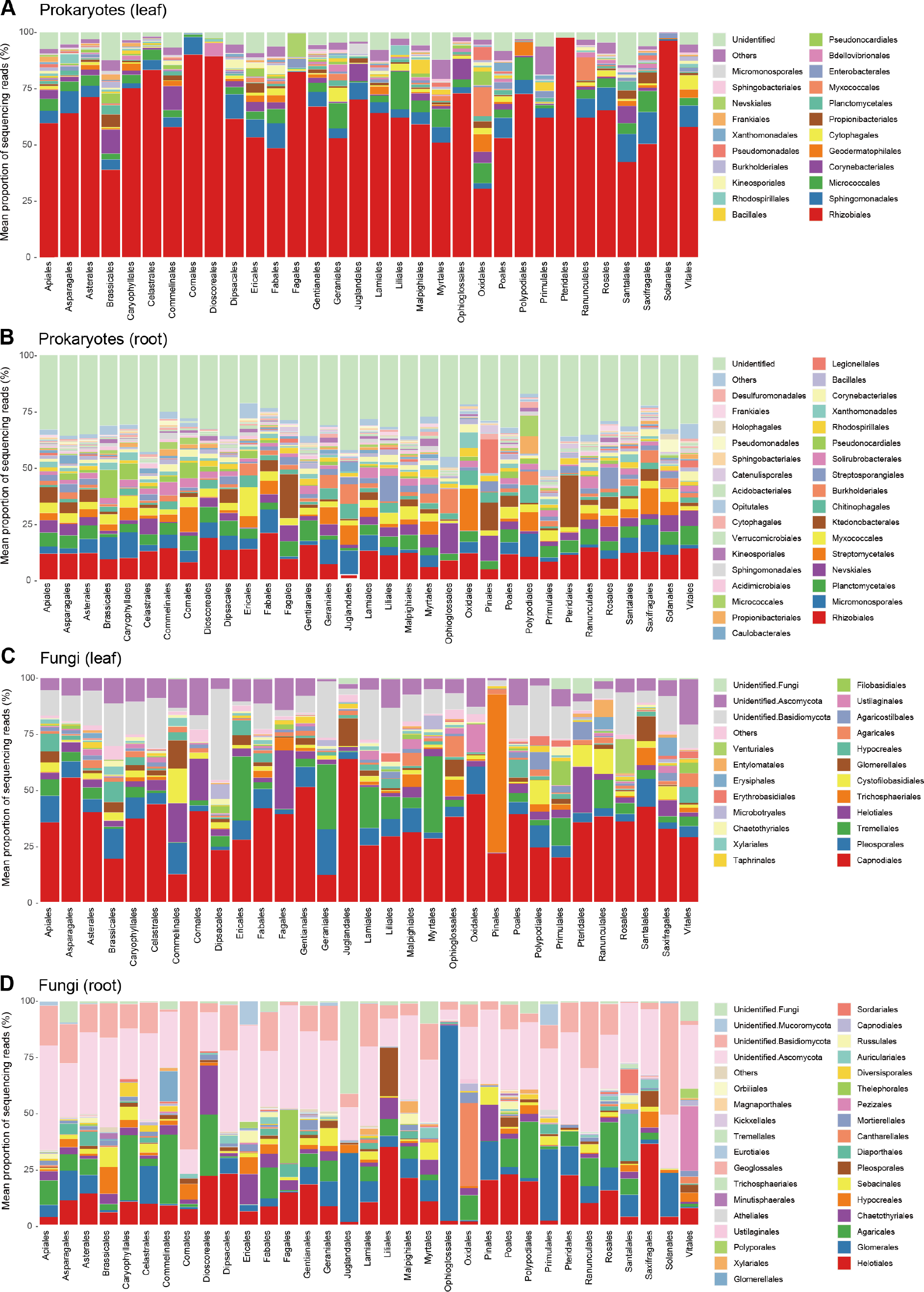
Prokaryote and fungal community compositions. **(A)** Order-level compositions of prokaryotes in leaf samples. Mean proportions of sequencing reads are shown for each plant order. In total, sequencing data were successfully obtained from 188 leaf samples. **(B)** Order-level compositions of prokaryotes in root samples (270 root samples). **(C)** Order-level compositions of fungi in leaf samples (213 leaf samples). Mean proportions of sequencing reads are shown for each plant order. **(D)** Order-level compositions of fungi in root samples (261 root samples).

### Factors Contributing to Microbiome Compositions

In the PERMANOVA, sampling month had significant effects on the leaf prokaryote, root prokaryote, and leaf fungal community compositions but not on the root fungal community structure (Table 1; Supplementary Table 1). Meanwhile, order-level host taxonomy influenced the root prokaryote, leaf fungal, and the root fungal community compositions but not the leaf prokaryote community structure (Table 1). The nativeness of host plants (native or alien) had significant impacts on the root prokaryote and the root fungal (genus-level) community compositions (Table 1). The analysis also showed that host plant lifeform (herbaceous or woody) had significant effects on the leaf fungal community structure (Table 1).

**TABLE 1.**
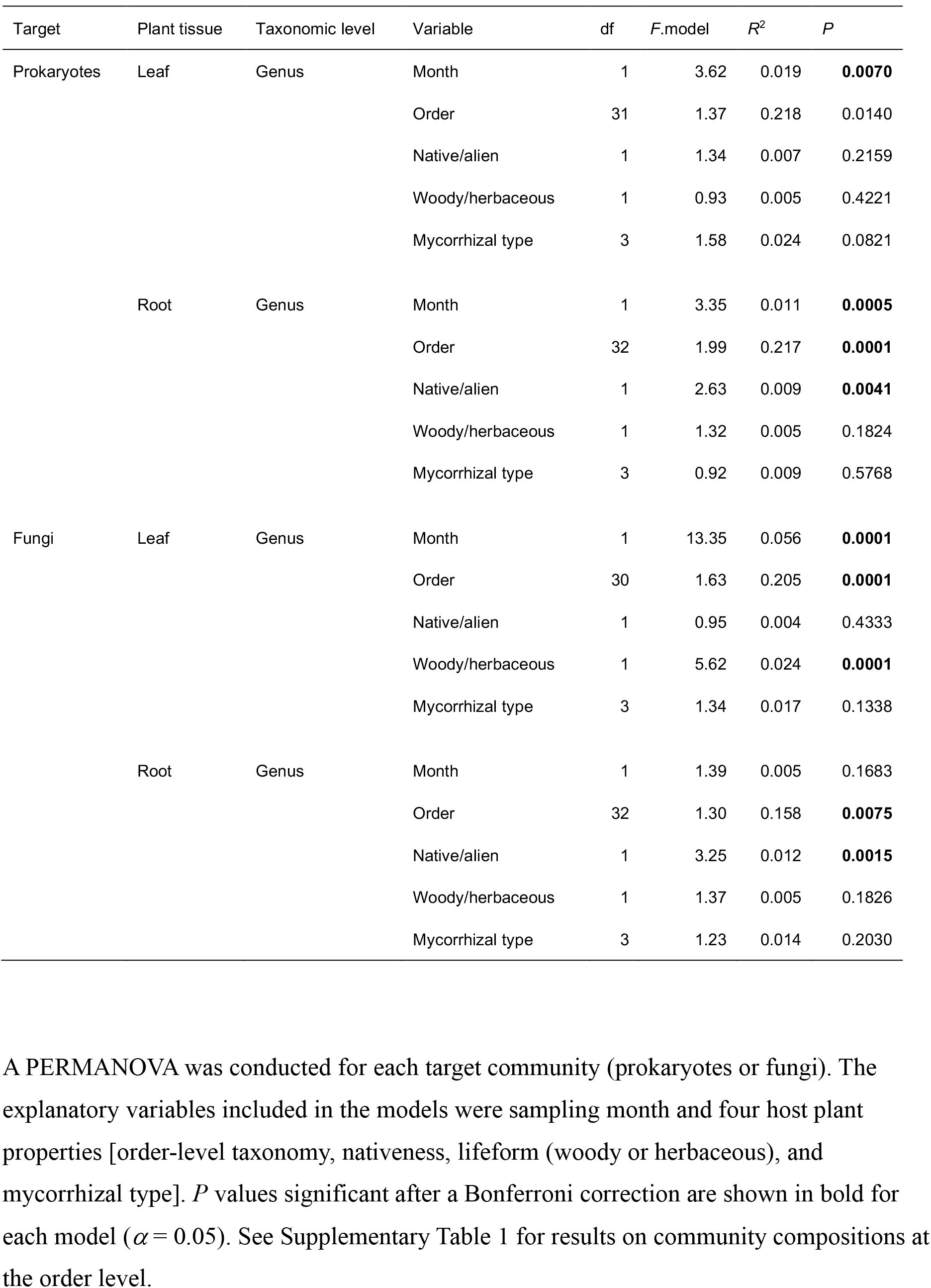
Factors contributing to variation in genus-level community compositions of bacteria and fungi.

| Target | Plant tissue | Taxonomic level | Variable | df | F.model | R<sup>2</sup> | P |
| --- | --- | --- | --- | --- | --- | --- | --- |
| Prokaryotes | Leaf | Genus | Month | 1 | 3.62 | 0.019 | **0.0070** |
|  |  |  | Order | 31 | 1.37 | 0.218 | 0.0140 |
|  |  |  | Native/alien | 1 | 1.34 | 0.007 | 0.2159 |
|  |  |  | Woody/herbaceous | 1 | 0.93 | 0.005 | 0.4221 |
|  |  |  | Mycorrhizal type | 3 | 1.58 | 0.024 | 0.0821 |
|  | Root | Genus | Month | 1 | 3.35 | 0.011 | **0.0005** |
|  |  |  | Order | 32 | 1.99 | 0.217 | **0.0001** |
|  |  |  | Native/alien | 1 | 2.63 | 0.009 | **0.0041** |
|  |  |  | Woody/herbaceous | 1 | 1.32 | 0.005 | 0.1824 |
|  |  |  | Mycorrhizal type | 3 | 0.92 | 0.009 | 0.5768 |
| Fungi | Leaf | Genus | Month | 1 | 13.35 | 0.056 | **0.0001** |
|  |  |  | Order | 30 | 1.63 | 0.205 | **0.0001** |
|  |  |  | Native/alien | 1 | 0.95 | 0.004 | 0.4333 |
|  |  |  | Woody/herbaceous | 1 | 5.62 | 0.024 | **0.0001** |
|  |  |  | Mycorrhizal type | 3 | 1.34 | 0.017 | 0.1338 |
|  | Root | Genus | Month | 1 | 1.39 | 0.005 | 0.1683 |
|  |  |  | Order | 32 | 1.30 | 0.158 | **0.0075** |
|  |  |  | Native/alien | 1 | 3.25 | 0.012 | **0.0015** |
|  |  |  | Woody/herbaceous | 1 | 1.37 | 0.005 | 0.1826 |
|  |  |  | Mycorrhizal type | 3 | 1.23 | 0.014 | 0.2030 |

### Randomization Analyses of Preferences

In the randomization analyses, the relative abundances of four bacterial and eight fungal genera changed through the sampling months (Table 2). For example, the fungal genera *Leucosporidium*, *Taphrina*, and *Dioszegia* in the leaf fungal community appeared preferentially in July, while the bacterial genera *Amnibacterium*, *Spirosoma*, and *Hymenobacter* preferentially occurred in September (Table 2). Regarding the nativeness of hosts, 14 bacterial and 23 fungal genera showed preferences for alien plant species (Table 3). The list of bacterial genera with preferences for alien plant species included *Deinococcus*, *Dermacoccus*, *Rubrobacter*, *Brevundimonas*, *Paraburkholderia*, and *Virgisporangium*, while that of fungal genera showing preferences for alien plants involved *Phoma*, *Hymenoscyphus*, *Sakaguchia*, *Didymella*, *Curvularia*, *Cylindrocarpon*, and *Meliniomyces* (Table 3). In contrast, two bacterial genera, *Actinoallomurus* and *Singulisphaera* showed preferences for native plant species (Table 3). The randomization analyses also indicated that two bacterial (*Massilia* and *Steroidobacter*) and four fungal (*Veronaea*, *Lophiostoma*, *Agrocybe*, and *Leptodontidium*) genera occurred preferentially on woody plant species (Supplementary Table 2). Although mycorrhizal type of host plants did not have significant effects in the community-level statistical analysis (Table 1), a number of bacterial and fungal genera showed preferences for host mycorrhizal type (Table 4; Supplementary Table 3). For example, the bacterial genera *Ferrimicrobium*, *Kineococcus*, *Sandarakinorhabdus*, and *Microthrix* showed preferences for non-mycorrhizal plant species, while *Flavisolibacter*, *Neochlamydia*, and *Phenylobacterium* showed preferences for ectomycorrhizal plants (Table 4). Fungi in the genera *Colletotrichum*, *Entorrhiza*, *Mycoarthris*, and *Sugiyamaella*, for instance, occurred preferentially on non-mycorrhizal plants, while not only ectomycorrhizal fungal genera (*Laccaria* and *Tomentella*) but also potentially endophytic fungal genera such as *Phialocephala* and *Oidiodendron* appeared preferentially on ectomycorrhizal plant species (Table 4; Supplementary Table 3). Bacteria and fungi with preferences for arbuscular mycorrhizal plants were not detected in the present randomization analyses presumably due to the dominance of arbuscular mycorrhizal plants within the datasets (see Discussion).

**TABLE 2.** Prokaryote and fungal genera showing preferences for months.

| Target | Plant tissue | Preferred month | Phylum | Genus | Preference score | FDR |
| --- | --- | --- | --- | --- | --- | --- |
| Prokaryotes | Leaf | July | Proteobacteria | *Bradyrhizobium* | 3.48 | 0.00948 |
|  |  | September | Actinobacteria | *Amnibacterium* | 3.60 | 0.00159 |
|  |  | September | Bacteroidetes | *Spirosoma* | 3.48 | 0.00211 |
|  |  | September | Bacteroidetes | *Hymenobacter* | 3.08 | 0.00948 |
| Fungi | Leaf | July | Basidiomycota | *Leucosporidium* | 4.32 | 0.00026 |
|  |  | July | Ascomycota | *Taphrina* | 3.93 | 0.00064 |
|  |  | July | Basidiomycota | *Dioszegia* | 3.11 | 0.01246 |
|  |  | August | Ascomycota | *Bryochiton* | 3.28 | 0.00058 |
|  |  | September | Ascomycota | *Nigrospora* | 3.58 | 0.00052 |
|  |  | September | Ascomycota | *Paraphaeosphaeria* | 3.57 | 0.00007 |
|  | Root | July | Basidiomycota | *Agrocybe* | 3.50 | 0.00384 |
|  |  | September | Ascomycota | *Paraphaeosphaeria* | 3.01 | 0.00686 |

Prokaryote and fungal genera are listed with their preference scores for specific months. The *P* values obtained in the randomization analysis were converted to false discovery rates (FDRs). Genera whose preference scores exceeded three are shown.

**TABLE 3.**
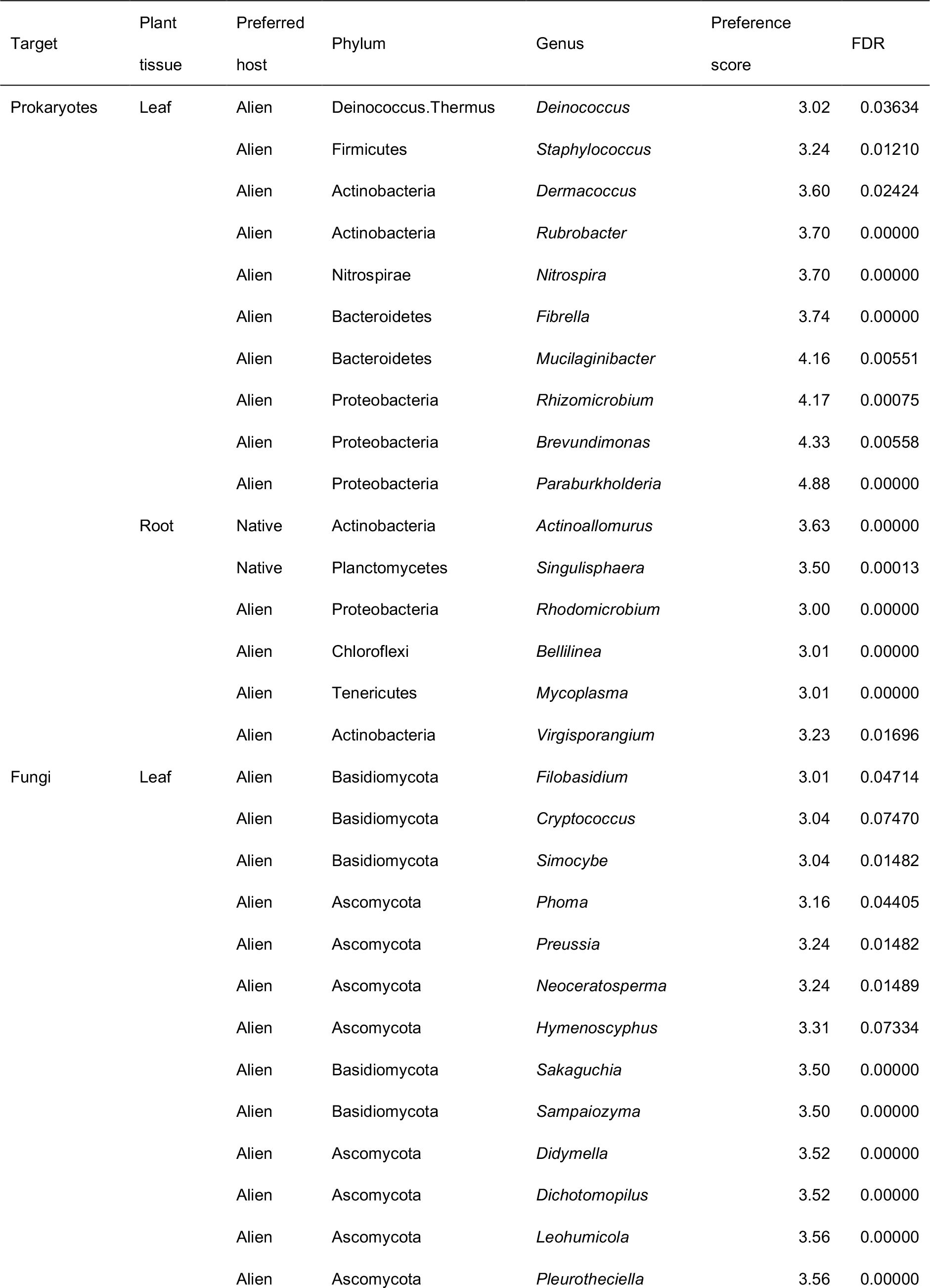
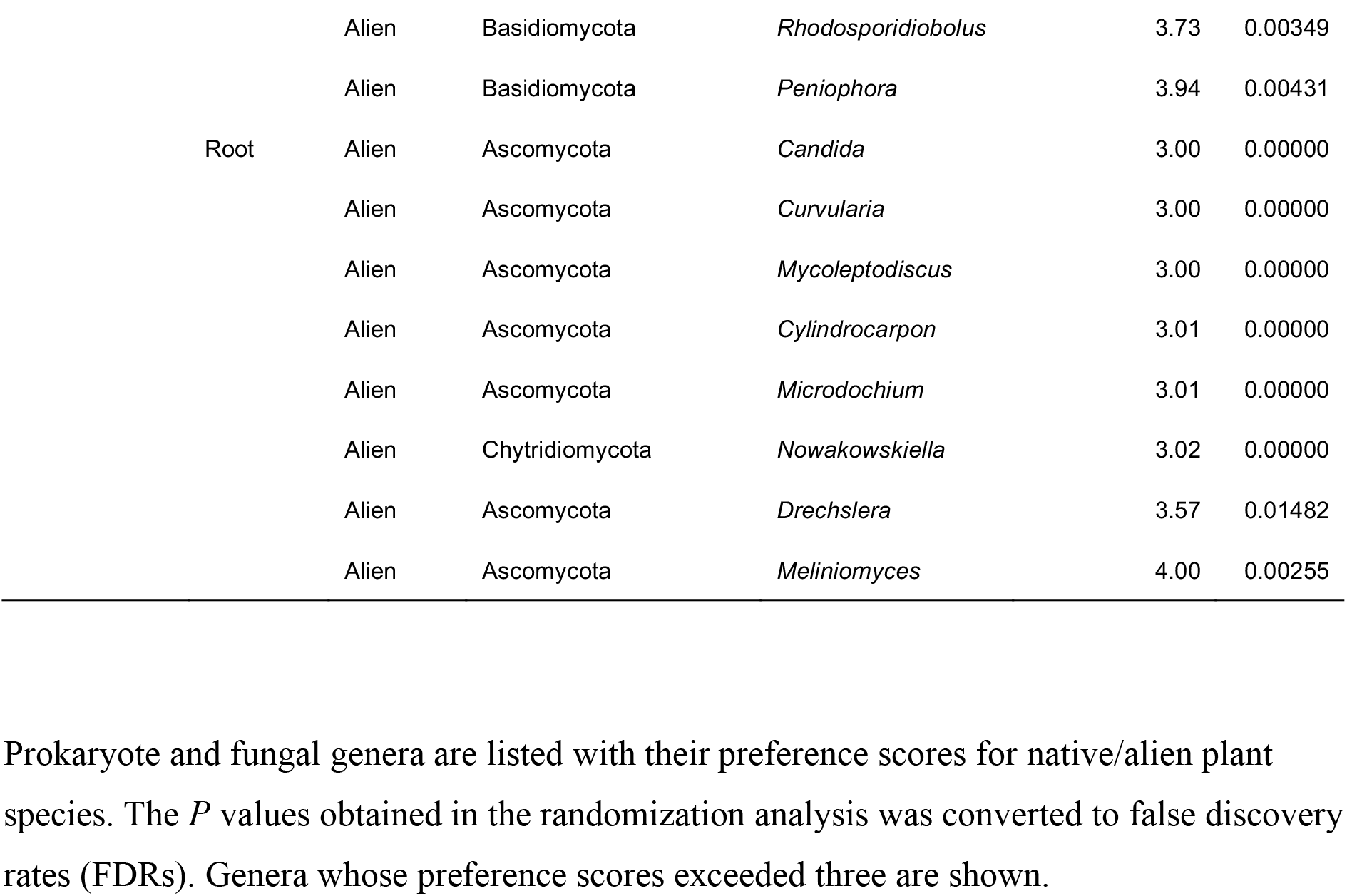
Prokaryote and fungal genera showing preferences for native/alien plant species.

**TABLE 4.**
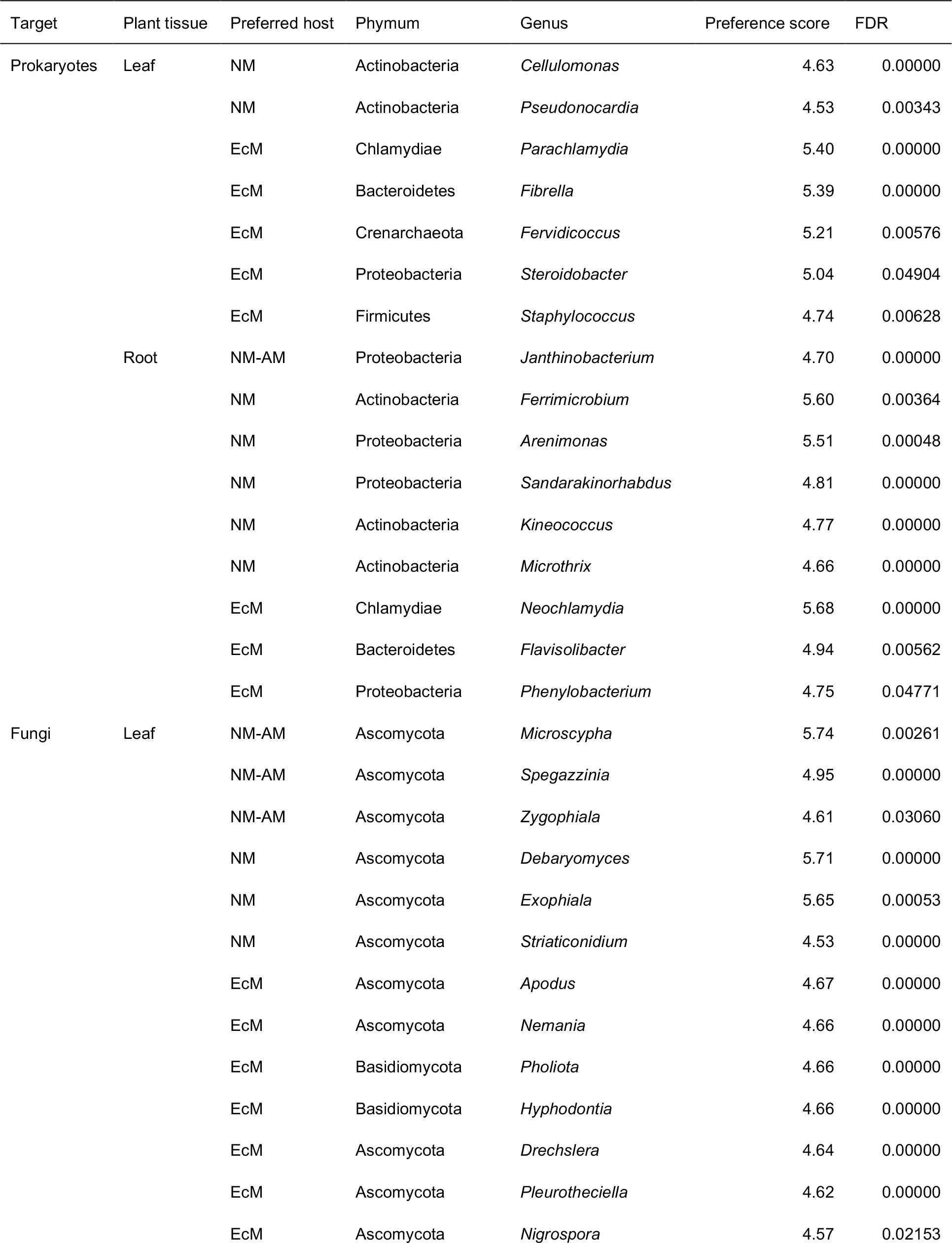
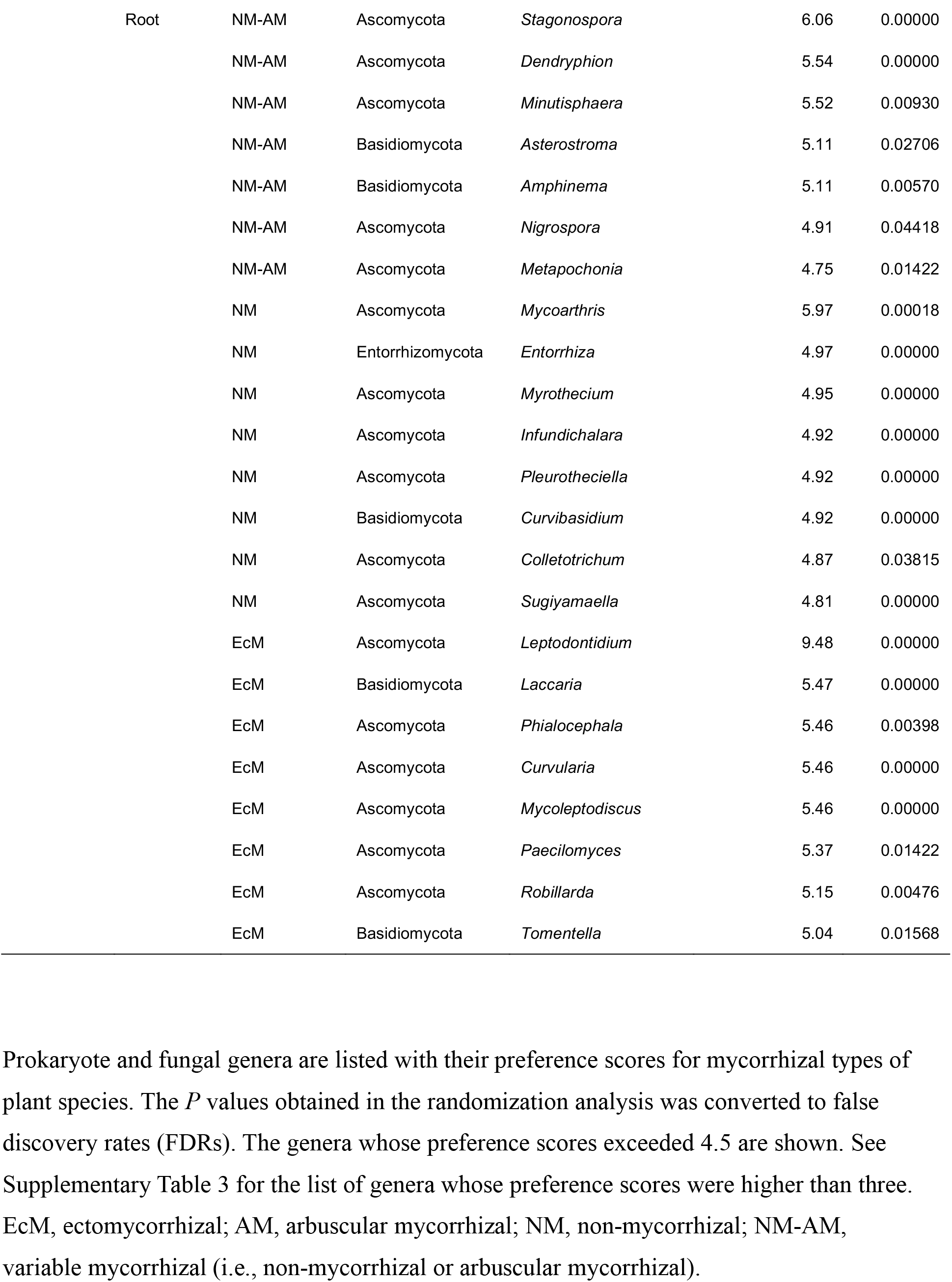
Prokaryote and fungal genera showing preferences for mycorrhizal types of host plant species.

## DISCUSSION

Based on a high-throughput sequencing dataset, we herein compared leaf and root microbiome compositions across co-occurring plant species in a temperate grassland. By targeting one of the most plant-species-rich ecosystems in the cool-temperate climate, we compared leaf- and root-associated microbial communities across 33 plant orders (Fig. 3) and then performed a series of statistical analyses on factors that may influence community compositions of plant-associated microbes (Tables 1-4). Hereafter, we discuss potential contributions of the factors examined, focusing on preferences of each microbial taxon for host characteristics.

An interesting finding of this study is that, while the compositions of leaf prokaryote, root prokaryote, and leaf fungal communities changed through the sampling months, root fungal community compositions did not significantly shift during the period (Tables 1-2). This pattern possibly represents difference in basic environmental features between above- and below-ground systems and/or difference in phenological patterns between prokaryote and fungal communities. For example, above-ground biotic/abiotic environments may be more dynamics than below-ground environments, resulting in rapid turnover of microbial communities. Moreover, above-ground parts of plants are more likely to be accessed by wind-dispersed spores and inocula than below-ground parts: hence, above-ground microbiome processes may be susceptible to continual immigration. In addition to potential contrasting features of above-vs. below-ground systems, difference in basic ecology between bacteria and fungi may have contributed to the varied phenological patterns. While mycorrhizal and endophytic fungi usually persist on/around host root systems in the form of hyphal networks (Lian et al., 2006;Smith and Read, 2008), bacterial communities may consist mainly of opportunistic symbionts [*sensu* (Hardoim et al., 2008)], which undergo rapid population growth under favorable environmental conditions and are subsequently replaced by others. Year-round comparative studies on leaf and root microbiomes are awaited for gaining more comprehensive understandings of microbiome dynamics.

Among the microbial communities examined, both root-associated prokaryote and fungal communities significantly varied between native and alien plant species (Table 1). The randomization analysis then allowed us to screen for bacterial and fungal genera showing preferences for native or alien plants (Table 3). Among the bacterial genera showing preferences for alien plants, *Paraburkholderia* has been known to include species with nitrogen-fixing abilities (Dall’Agnol et al., 2016), potentially influencing host nutritional conditions. In addition, the analysis showed that various genera in the phylum Actinobacteria (*Rubrobacter*, *Dermacoccus*, *Actinoallomurus*, and *Virgisporangium*) showed preferences for native or alien plant species. Given that many actinomycete bacteria produce chemicals suppressing other microbes (Qin et al., 2011;Bérdy, 2012), their ecological roles in ecosystems are of particular interest. For fungi, although the absence of OTUs displaying preferences for native plants requires careful interpretation (see below), the randomization analysis showed that various fungal taxa could have preferences for alien plant species (Table 3). Among them, *Curvularia*, *Didymella*, and *Cylindrocarpon* include well-characterized plant pathogenic species (Alaniz et al., 2007;Akinbode, 2010;Keinath, 2011). In contrast, fungi in the genus *Meliniomyces* have been described as mycorrhizal or endophytic fungi (Hambleton and Sigler, 2005;Ohtaka and Narisawa, 2008;Vohník et al., 2013), possibly contributing to the survival and growth of host plants. Overall, these results suggest that various taxonomic groups of bacteria and fungi are associated with native or alien plant species, potentially affecting invasiveness of alien plants both positively and negatively.

At the whole community level, mycorrhizal types of host plants did not have significant effects on plant microbiome compositions, while effects of plant lifeform (herbaceous or woody) were significant in one of the communities examined (i.e., leaf fungal community) (Table 1; Supplementary Table 3). However, the randomization analysis for respective microbial taxa highlighted diverse bacterial and fungal genera showing statistically significant preferences for host mycorrhizal types (Table 4). Among the bacteria showing preferences for non-mycorrhizal plants, *Ferrimicrobium* includes species adapted to low pH conditions (Johnson et al., 2009), while the genus *Kineococcus* is known to involve species tolerant to salt stress (Bian et al., 2012). Within the fungal community, ectomycorrhizal fungi in the genera *Laccaria* and *Tomentella* showed preferences for ectomycorrhizal plant species as expected based on previous studies on mycorrhizal symbioses (Smith and Read, 2008;Tedersoo et al., 2010). We also found that possibly endophytic fungi in the genus *Phialocephala* (Fernando and Currah, 1996;Grünig et al., 2008) showed preferences for ectomycorrhizal plants. Among the fungal genera showing preferences for non-mycorrhizal plant roots, *Colletotrichum* is of particular interest. Although many *Colletotrichum* species had been known as plant pathogens (Hammerschmidt et al., 1982;O’Connell et al., 2012), recent studies have demonstrated that some species in the genus could work as mutualistic symbionts by providing soil phosphorus to non-mycorrhizal plants such as *Arabidopsis thaliana* (Hiruma et al., 2016;Hiruma et al., 2018). Thus, the list of microbes preferentially associated with non-mycorrhizal plants (Table 4; Supplementary Table 3) sheds light on potential diversity of bacteria and fungi that may partly fill niches of mycorrhizal fungi in non-mycorrhizal plant species.

Although the data collected in this study provide fundamental information of microbial diversity in a grassland ecosystem, the statistical results should be interpreted with caution. First, the small number of samples per plant species may have affected the comparison of microbiome compositions among plant taxa (Fig. 3). The identification of plant roots is time-consuming especially in species-rich grasslands consisting mainly of perennial plants with tangled root systems, limiting the throughput of sampling. Therefore, for more comprehensive profiling of plant microbiomes, we may need to increase the throughput of plant species identification based on molecular taxonomic assignment (i.e., DNA barcoding) of host plants (Hollingsworth et al., 2009;Toju et al., 2013). Second, the presence of unidentified bacteria and fungi in the dataset may have biased the statistical analyses. Although databases of microbes have been continually updated, there remain many bacterial and fungal lineages whose taxonomy has not yet been fixed. In particular, below-ground microbiomes are known to involve a number of poorly investigated taxa, whose physiological and ecological functions remain to be uncovered (Buée et al., 2009;Fierer, 2017). Thus, with more reference microbial databases, we will be able to examine whether the patterns found in the present analysis hold after assigning unidentified OTUs to right categories. Third, there seems limitation of the randomization method used in this study. In the analysis of host plant nativeness, significant preferences for native plant species were detected only for a few microbial genera (Table 3). Likewise, in the analysis on plant mycorrhizal types, there was no microbial genus showing preferences for arbuscular mycorrhizal plants (Table 4). Given that native and arbuscular mycorrhizal plant species were dominant in the grassland [87.9 % (254/289) and 87.5 % (253/289) of samples, respectively], this kind of randomization analyses may tend to provide conservative results for major categories, while yielding sensitive results for categories with smaller number of samples. Although randomization methods require fewer statistical assumptions than model-based methods [e.g., (Sato et al., 2015)], they may be more suitable for data matrices with equal number of replicate samples across target categories.

We herein revealed how diverse bacterial and fungal taxa were associated with leaves and roots of the 138 plant species co-occurring in a cool-temperate grassland, focusing on potential contributions of host plant characteristics on microbiome compositions. Although recent ecological studies have highlighted possible feedbacks between plant and microbial community dynamics (Bever et al., 2010;Mangan et al., 2010;Van der Putten et al., 2013), we still have limited knowledge of the processes by which species-rich plant communities are maintained by phyllosphere and rhizosphere microbiomes. Accumulating comprehensive inventory data of microbiomes associated with whole plant communities is a prerequisite for advancing our understanding of ecosystem-scale processes. Case studies in various types of terrestrial ecosystems in diverse climatic regions will allow us to elucidate how plant species with different mycorrhizal types often coexist in natural ecosystems (Booth, 2004;Kadowaki et al., in press) or why some ecosystems are resistant against alien plants, while others are heavily disturbed by invasive species (Mitchell and Power, 2003;Reinhart and Callaway, 2006).

## AUTHOR CONTRIBUTIONS

HT, HK, and TK designed the work. HT, HK, and TK performed fieldwork. HT conducted molecular experiment and analyzed the data. HT, HK, and TK wrote the manuscript.

## ACKNOWLEDGEMENTS

We thank Qing-Wei Wang, Yuka Yamamoto, and Rie Watanabe for their help in fieldwork and Sarasa Amma and Hiroki Kawai for their support in molecular experiments. This work was financially supported by JST PRESTO (JPMJPR16Q6) to HT. This work was financially supported by JST PRESTO (JPMJPR16Q6) to HT, by the JSPS Grant-in-Aid for Scientific Research (B) Grant number JP17H03736 to HK, and by the promotion cost for functional enhancement of the Mountain Science Center to TK.

## SUPPLEEMENTARY MATERIAL

The Supplementary Material for this article can be found online at XXXXX.

## REFERENCES

Agler, M.T., Ruhe, J., Kroll, S., Morhenn, C., Kim, S.-T., Weigel, D., et al. (2016). Microbial hub taxa link host and abiotic factors to plant microbiome variation. PLOS Biol. 14, e1002352. doi: 10.1371/journal.pbio.1002352

Akinbode, O. (2010). Evaluation of antifungal efficacy of some plant extracts on Curvularia lunata, the causal organism of maize leaf spot. African J. ENc. Sci. Tech. 4, 797–800. doi:

Alaniz, S., León, M., Vicent, A., García-Jiménez, J., Abad-Campos, P., and Armengol, J. (2007). Characterization of Cylindrocarpon species associated with black foot disease of grapevine in Spain. Plant Disease 91, 1187–1193. doi:

Anderson, M.J. (2001). A new method for non-parametric multivariate analysis of variance. Austral Ecol. 26, 32–46. doi: 10.1111/j.1442-9993.2001.01070.pp.x

Apprill, A., Mcnally, S., Parsons, R., and Weber, L. (2015). Minor revision to V4 region SSU rRNA 806R gene primer greatly increases detection of SAR11 bacterioplankton. doi: 10.3354/ame01753

Arnold, A.E., Mejía, L.C., Kyllo, D., Rojas, E.I., Maynard, Z., Robbins, N., et al. (2003). Fungal endophytes limit pathogen damage in a tropical tree. Proc. Natl. Acad. Sci. USA. 100, 15649–15654. doi: 10.1073/pnas.2533483100

Bai, Y., Müller, D.B., Srinivas, G., Garrido-Oter, R., Potthoff, E., Rott, M., et al. (2015). Functional overlap of the Arabidopsis leaf and root microbiota. Nature 528, 364–369. doi: 10.1038/nature16192

Bérdy, J. (2012). Thoughts and facts about antibiotics: where we are now and where we are heading. J. Antibiotics 65, 385. doi:

Berendsen, R.L., Pieterse, C.M., and Bakker, P.A. (2012). The rhizosphere microbiome and plant health. Trends Plant Sci. 17, 478–486. doi: 10.1016/j.tplants.2012.04.001

Bever, J.D., Dickie, I.A., Facelli, E., Facelli, J.M., Klironomos, J., Moora, M., et al. (2010). Rooting theories of plant community ecology in microbial interactions. Trends Ecol. Evol. 25, 468–478. doi: 10.1016/j.tree.2010.05.004

Bian, G.-K., Feng, Z.-Z., Qin, S., Xing, K., Wang, Z., Cao, C.-L., et al. (2012). Kineococcus endophytica sp. nov., a novel endophytic actinomycete isolated from a coastal halophyte in Jiangsu, China. Antonie van Leeuwenhoek 102, 621–628. doi:

Booth, M.G. (2004). Mycorrhizal networks mediate overstorey-understorey competition in a temperate forest. Ecol. Lett. 7, 538–546. doi:

Brundrett, M.C. (2009). Mycorrhizal associations and other means of nutrition of vascular plants: understanding the global diversity of host plants by resolving conflicting information and developing reliable means of diagnosis. Plant Soil 320, 37–77. doi: 10.1007/s11104-008-9877-9

Buée, M., Reich, M., Murat, C., Morin, E., Nilsson, R., Uroz, S., et al. (2009). 454 Pyrosequencing analyses of forest soils reveal an unexpectedly high fungal diversity. New Phytol. 184, 449–456. doi: 10.1111/j.1469-8137.2009.03003.x

Caporaso, J.G., Lauber, C.L., Walters, W.A., Berg-Lyons, D., Lozupone, C.A., Turnbaugh, P.J., et al. (2011). Global patterns of 16S rRNA diversity at a depth of millions of sequences per sample. Proc. Natl. Acad. Sci. USA. 108, 4516–4522. doi: 10.1073/pnas.1000080107

Compant, S., Duffy, B., Nowak, J., Clément, C., and Barka, E.A. (2005). Use of plant growth-promoting bacteria for biocontrol of plant diseases: principles, mechanisms of action, and future prospects. Appl. Env. Microbiol. 71, 4951–4959. doi:

Dall’agnol, R.F., Plotegher, F., Souza, R.C., Mendes, I.C., Dos Reis Junior, F.B., Béna, G., et al. (2016). Paraburkholderia nodosa is the main N2-fixing species trapped by promiscuous common bean (Phaseolus vulgaris L.) in the Brazilian ‘Cerradão’. FEMS Microbiol. Ecol. 92, fiw108. doi:

Delmotte, N., Knief, C., Chaffron, S., Innerebner, G., Roschitzki, B., Schlapbach, R., et al. (2009). Community proteogenomics reveals insights into the physiology of phyllosphere bacteria. Proc. Natl. Acad. Sci. USA 106, 16428–16433. doi:

Fernando, A.A., and Currah, R.S. (1996). A comparative study of the effects of the root endophytes Leptodontidium orchidicola and Phialocephala fortinii (Fungi Imperfecti) on the growth of some subalpine plants in culture. Can. J. Bot. 74, 1071–1078. doi:

Fierer, N. (2017). Embracing the unknown: disentangling the complexities of the soil microbiome. Nat. Rev. Microbiol. 15, 579–590. doi:

Frey-Klett, P., Garbaye, J.A., and Tarkka, M. (2007). The mycorrhiza helper bacteria revisited. New Phytol. 176, 22–36. doi:

Gao, F.-K., Dai, C.-C., and Liu, X.-Z. (2010). Mechanisms of fungal endophytes in plant protection against pathogens. African J. Microbiol. Res. 4, 1346–1351. doi:

Grünig, C.R., Queloz, V., Sieber, T.N., and Holdenrieder, O. (2008). Dark septate endophytes (DSE) of the Phialocephala fortinii s.1. – Acephala applanata pecies complex in tree roots: classification, population biology, and ecology. Botany 86, 1355–1369. doi:

Hacquard, S., Garrido-Oter, R., González, A., Spaepen, S., Ackermann, G., Lebeis, S., et al. (2015). Microbiota and host nutrition across plant and animal kingdoms. Cell Host Microbe 17, 603–616. doi:

Hacquard, S., Spaepen, S., Garrido-Oter, R., and Schulze-Lefert, P. (2017). Interplay between innate immunity and the plant microbiota. Ann. Rev. Phytopathol. 55, 565–589. doi:

Hamady, M., Walker, J.J., Harris, J.K., Gold, N.J., and Knight, R. (2008). Error-correcting barcoded primers for pyrosequencing hundreds of samples in multiplex. Nat. Methods 5, 235–237. doi: 10.1038/nmeth.1184

Hambleton, S., and Sigler, L. (2005). Meliniomyces, a new anamorph genus for root-associated fungi with phylogenetic affinities to Rhizoscyphus ericae(≡ Hymenoscyphus ericae), Leotiomycetes. Studies Mycology 53, 1–27. doi:

Hammerschmidt, R., Nuckles, E., and Kuć, J. (1982). Association of enhanced peroxidase activity with induced systemic resistance of cucumber to Colletotrichum lagenarium. Phisiol. Plant Pathol. 20, 73–82. doi:

Hardoim, P.R., Van Overbeek, L.S., and Van Elsas, J.D. (2008). Properties of bacterial endophytes and their proposed role in plant growth. Trends Microbiol. 16, 463–471. doi:

Hiruma, K., Gerlach, N., Sacristán, S., Nakano, R.T., Hacquard, S., Kracher, B., et al. (2016). Root endophyte Colletotrichum tofieldiae confers plant fitness benefits that are phosphate status dependent. Cell 165, 464–474. doi:

Hiruma, K., Kobae, Y., and Toju, H. (2018). Beneficial associations between Brassicaceae plants and fungal endophytes under nutrient-limiting conditions: evolutionary origins and host–symbiont molecular mechanisms. Current Opiion Plant Biol. 44, 145–154. doi: 10.1016/j.pbi.2018.04.009

Hoffman, M.T., and Arnold, A.E. (2010). Diverse bacteria inhabit living hyphae of phylogenetically diverse fungal endophytes. Appl. Env. Microbiol. 76, 4063–4075. doi:

Hollingsworth, P.M., Forrest, L.L., Spouge, J.L., Hajibabaei, M., Ratnasingham, S., Van Der Bank, M., et al. (2009). A DNA barcode for land plants. Proc. Natl. Acad. Sci. USA 106, 12794–12797. doi: 10.1073/pnas.0905845106

Huson, D.H., Auch, A.F., Qi, J., and Schuster, S.C. (2007). MEGAN analysis of metagenomic data. Genome Res. 17, 377–386. doi: 10.1101/gr.5969107

Johnson, D.B., Bacelar-Nicolau, P., Okibe, N., Thomas, A., and Hallberg, K.B. (2009). Ferrimicrobium acidiphilum gen. nov., sp. nov. and Ferrithrix thermotolerans gen. nov., sp. nov.: heterotrophic, iron-oxidizing, extremely acidophilic actinobacteria. Int. J. Syst. Evol. Microbiol. 59, 1082–1089. doi:

Kadowaki, K., Yamamoto, S., Sato, H., Tanabe, A.S., Hidaka, A., and Toju, H. (in press). Mycorrhizal fungi mediate the direction and strength of plant-soil feedbacks differently between arbuscular mycorrhizal and ectomycorrhizal communities. Commun. Biol. doi:

Keinath, A.P. (2011). From native plants in central Europe to cultivated crops worldwide: the emergence of Didymella bryoniae as a cucurbit pathogen. HortScience 46, 532–535. doi:

Lian, C., Narimatsu, M., Nara, K., and Hogetsu, T. (2006). Tricholoma matsutake in a natural Pinus densiflora forest: correspondence between above-and below-ground genets, association with multiple host trees and alteration of existing ectomycorrhizal communities. New Phytol. 171, 825–836. doi:

Lundberg, D.S., Lebeis, S.L., Paredes, S.H., Yourstone, S., Gehring, J., Malfatti, S., et al. (2012). Defining the core Arabidopsis thaliana root microbiome. Nature 488, 86–90. doi: 10.1038/nature11237

Lundberg, D.S., Yourstone, S., Mieczkowski, P., Jones, C.D., and Dangl, J.L. (2013). Practical innovations for high-throughput amplicon sequencing. Nat. Methods 10, 999–1002. doi: 10.1038/nmeth.2634

Mangan, S.A., Schnitzer, S.A., Herre, E.A., Mack, K.M., Valencia, M.C., Sanchez, E.I., et al. (2010). Negative plant-soil feedback predicts tree-species relative abundance in a tropical forest. Nature 466, 752–755. doi: 10.1038/nature09273

Mercier, J., and Lindow, S. (2000). Role of leaf surface sugars in colonization of plants by bacterial epiphytes. Appl. Env. Microbiol. 66, 369–374. doi:

Mitchell, C.E., and Power, A.G. (2003). Release of invasive plants from fungal and viral pathogens. Nature 421, 625. doi:

Newsham, K.K. (2011). A meta-analysis of plant responses to dark septate root endophytes. New Phytol. 190, 783–793. doi: 10.1111/j.1469-8137.2010.03611.x

O’connell, R.J., Thon, M.R., Hacquard, S., Amyotte, S.G., Kleemann, J., Torres, M.F., et al. (2012). Lifestyle transitions in plant pathogenic Colletotrichum fungi deciphered by genome and transcriptome analyses. Nat. Genetics 44, 1060. doi:

Ohtaka, N., and Narisawa, K. (2008). Molecular characterization and endophytic nature of the root-associated fungus Meliniomyces variabilis (LtVB3). J. General Plant Pathol. 74, 24–31. doi:

Oksanen, J., Blanachet, F.G., Kindt, R., Legendre, P., Minchin, P.R., O’hara, R.B., et al. (2012). “Vegan: community ecology package. R package version 2.0-3 available at http://CRAN.R-project.org/package=vegan”.).

Öpik, M., Metsis, M., Daniell, T., Zobel, M., and Moora, M. (2009). Large-scale parallel 454 sequencing reveals host ecological group specificity of arbuscular mycorrhizal fungi in a boreonemoral forest. New Phytol. 184, 424–437. doi: 10.1111/j.1469-8137.2009.02920.x

Parniske, M. (2008). Arbuscular mycorrhiza: the mother of plant root endosymbioses. Nat. Rev. Microbiol. 6, 763–775. doi:

Peay, K.G., Kennedy, P.G., and Talbot, J.M. (2016). Dimensions of biodiversity in the Earth mycobiome. Nat. Rev. Microbiol. 14, 434–447. doi: 10.1038/nrmicro.2016.59

Peay, K.G., Russo, S.E., Mcguire, K.L., Lim, Z., Chan, J.P., Tan, S., et al. (2015). Lack of host specificity leads to independent assortment of dipterocarps and ectomycorrhizal fungi across a soil fertility gradient. Ecol. Lett. 18, 807–816. doi: 10.1111/ele.12459

Pieterse, C.M., Zamioudis, C., Berendsen, R.L., Weller, D.M., Van Wees, S.C., and Bakker, P.A. (2014). Induced systemic resistance by beneficial microbes. Ann. Rev. Phytopathol. 52, 347–375. doi:

Qin, S., Xing, K., Jiang, J.-H., Xu, L.-H., and Li, W.-J. (2011). Biodiversity, bioactive natural products and biotechnological potential of plant-associated endophytic actinobacteria. Appl. Microbiol. Biotech. 89, 457–473. doi:

R-Core-Team (2018). “R 3.5.1: A language and environment for statistical computing available at http://www.R-project.org/”. (Vienna, Austri: R Foundation for Statistical Computing).

Ramamoorthy, V., Viswanathan, R., Raguchander, T., Prakasam, V., and Samiyappan, R. (2001). Induction of systemic resistance by plant growth promoting rhizobacteria in crop plants against pests and diseases. Crop Protection 20, 1–11. doi:

Reinhart, K.O., and Callaway, R.M. (2006). Soil biota and invasive plants. New Phytol. 170, 445–457. doi:

Rognes, T., Mahé, F., Flouri, T., Quince, C., and Nichols, B. (2014). “Vsearch: program available at https://github.com/torognes/vsearch.”.).

Sato, H., Tanabe, A.S., and Toju, H. (2015). Contrasting diversity and host association of ectomycorrhizal Basidiomycetes versus root-associated Ascomycetes in a dipterocarp rainforest. PLOS ONE 10, e0125550. doi: 10.1371/journal.pone.0125550

Schardl, C.L., and Phillips, T.D. (1997). Protective grass endophytes: where are they from and where are they going? Plant Disease 81, 430–438. doi:

Schlaeppi, K., and Bulgarelli, D. (2015). The plant microbiome at work. Mol. Plant-Microbe Int. 28, 212–217. doi: 10.1094/MPMI-10-14-0334-FI

Smith, S.E., and Read, D.J. (2008). Mycorrhizal symbiosis. New York: Academic press.

Stevens, J.L., Jackson, R.L., and Olson, J.B. (2013). Slowing PCR ramp speed reduces chimera formation from environmental samples. J. Microbiol. Methods 93, 203–205. doi: 10.1016/j.mimet.2013.03.013

Tanabe, A. S. (2018). Claident v0.2.2018.05.29, a software distributed by author at http://www.fifthdimension.jp/.

Tanabe, A.S., and Toju, H. (2013). Two new computational methods for universal DNA barcoding: a benchmark using barcode sequences of bacteria, archaea, animals, fungi, and land plants. PLOS ONE 8, e76910. doi: 10.1371/journal.pone.0076910

Tedersoo, L., May, T.W., and Smith, M.E. (2010). Ectomycorrhizal lifestyle in fungi: global diversity, distribution, and evolution of phylogenetic lineages. Mycorrhiza 20, 217–263. doi:

Toju, H., Guimarães, P.R., Jr, Olesen, J.M., and Thompson, J.N. (2014). Assembly of complex plant–fungus networks. Nat. Commun. 5, 5273. doi: 10.1038/ncomms6273

Toju, H., Peay, K.G., Yamamichi, M., Narisawa, K., Hiruma, K., Naito, K., et al. (2018a). Core microbiomes for sustainable agroecosystems. Nat. Plants 4, 247–257. doi: 10.1038/s41477-018-0139-4

Toju, H., Tanabe, A., and Ishii, H. (2016a). Ericaceous plant–fungus network in a harsh alpine–subalpine environment. Mol. Ecol. 25, 3242–3257. doi: 10.1111/mec.13680

Toju, H., Tanabe, A.S., and Sato, H. (2018b). Network hubs in root-associated fungal metacommunities. Microbiome 6, 116. doi:

Toju, H., Tanabe, A.S., Yamamoto, S., and Sato, H. (2012). High-coverage ITS primers for the DNA-based identification of ascomycetes and basidiomycetes in environmental samples. PLOS ONE 7, e40863. doi: 10.1371/journal.pone.0040863

Toju, H., Yamamoto, S., Sato, H., Tanabe, A.S., Gilbert, G.S., and Kadowaki, K. (2013). Community composition of root-associated fungi in a Quercus-dominated temperate forest: “codominance” of mycorrhizal and root-endophytic fungi. Ecol. Evol. 3, 1281–1293. doi: 10.1002/ece3.546

Toju, H., Yamamoto, S., Tanabe, A.S., Hayakawa, T., and Ishii, H.S. (2016b). Network modules and hubs in plant-root fungal biome. J. R. Soc. Interface 13, 20151097. doi: 10.1098/rsif.2015.1097

Van Der Heijden, M.G., Bardgett, R.D., and Van Straalen, N.M. (2008). The unseen majority: soil microbes as drivers of plant diversity and productivity in terrestrial ecosystems. Ecol. Lett. 11, 296–310. doi:

Van Der Heijden, M.G., Klironomos, J.N., Ursic, M., Moutoglis, P., Streitwolf-Engel, R., Boller, T., et al. (1998). Mycorrhizal fungal diversity determines plant biodiversity, ecosystem variability and productivity. Nature 396, 69–72. doi:

Van Der Putten, W.H., Bardgett, R.D., Bever, J.D., Bezemer, T.M., Casper, B.B., Fukami, T., et al. (2013). Plant–soil feedbacks: the past, the present and future challenges. J. Ecol. 101, 265–276. doi:

Vohník, M., Mrnka, L., Lukešová, T., Bruzone, M.C., Kohout, P., and Fehrer, J. (2013). The cultivable endophytic community of Norway spruce ectomycorrhizas from microhabitats lacking ericaceous hosts is dominated by ericoid mycorrhizal Meliniomyces variabilis. Fungal Ecol. 6, 281–292. doi:

Wagner, M.R., Lundberg, D.S., Tijana, G., Tringe, S.G., Dangl, J.L., and Mitchell-Olds, T. (2016). Host genotype and age shape the leaf and root microbiomes of a wild perennial plant. Nature Commun. 7, 12151. doi:

